# Comprehensive single cell transcriptional profiling of a multicellular organism by combinatorial indexing

**DOI:** 10.1101/104844

**Authors:** Junyue Cao, Jonathan S. Packer, Vijay Ramani, Darren A. Cusanovich, Chau Huynh, Riza Daza, Xiaojie Qiu, Choli Lee, Scott N. Furlan, Frank J. Steemers, Andrew Adey, Robert H. Waterston, Cole Trapnell, Jay Shendure

**Affiliations:** Department of Genome Sciences, University of Washington, Seattle, WA, USA; Molecular and Cellular Biology Program, University of Washington, Seattle, WA, USA; Ben Towne Center for Childhood Cancer Research, Seattle Children’s Research Institute, Seattle, WA, USA; Department of Pediatrics, University of Washington, Seattle, WA, USA; Fred Hutchinson Cancer Research Center, Seattle, WA, USA; Illumina Inc., Advanced Research Group, San Diego, CA, USA; Department of Molecular & Medical Genetics, Oregon Health & Science University, Portland, OR, USA; Knight Cardiovascular Institute, Portland, OR, USA; Howard Hughes Medical Institute, Seattle, WA, USA

## Abstract

Conventional methods for profiling the molecular content of biological samples fail to resolve heterogeneity that is present at the level of single cells. In the past few years, single cell RNA sequencing has emerged as a powerful strategy for overcoming this challenge. However, its adoption has been limited by a paucity of methods that are at once simple to implement and cost effective to scale massively. Here, we describe a combinatorial indexing strategy to profile the transcriptomes of large numbers of single cells or single nuclei without requiring the physical isolation of each cell (<u>S</u>ingle cell <u>C</u>ombinatorial <u>I</u>ndexing RNA-seq or sci-RNA-seq). We show that sci-RNA-seq can be used to efficiently profile the transcriptomes of tens-of-thousands of single cells per experiment, and demonstrate that we can stratify cell types from these data. Key advantages of sci-RNA-seq over contemporary alternatives such as droplet-based single cell RNA-seq include sublinear cost scaling, a reliance on widely available reagents and equipment, the ability to concurrently process many samples within a single workflow, compatibility with methanol fixation of cells, cell capture based on DNA content rather than cell size, and the flexibility to profile either cells or nuclei. As a demonstration of sci-RNA-seq, we profile the transcriptomes of 42,035 single cells from *C. elegans* at the L2 stage, effectively 50-fold “shotgun cellular coverage” of the somatic cell composition of this organism at this stage. We identify 27 distinct cell types, including rare cell types such as the two distal tip cells of the developing gonad, estimate consensus expression profiles and define cell-type specific and selective genes. Given that *C. elegans* is the only organism with a fully mapped cellular lineage, these data represent a rich resource for future methods aimed at defining cell types and states. They will advance our understanding of developmental biology, and constitute a major step towards a comprehensive, single-cell molecular atlas of a whole animal.

## Introduction

Individual cells are the natural unit of form and function in biological systems. However, conventional methods for profiling the molecular content of biological samples usually mask cellular heterogeneity that is likely present even in ostensibly homogenous tissues. The averaging of signals from large numbers of single cells sharply limits what we are able to learn from the resulting data (*1*). For example, differences between samples cannot easily be attributed to differences within cells of the same type vs. differences in cell type composition.

In the past few years, profiling the transcriptome of individual cells (*i.e.* single cell RNA-seq) has emerged as a powerful strategy for resolving heterogeneity in biological samples. The expression levels of mRNA species are readily linked to cellular function, and therefore can be used to classify cell types in heterogeneous samples (*2–10*) as well as to order cell states in dynamic systems (*11*). Although methods for single cell RNA-seq have proliferated, they universally rely on the isolation of individual cells within physical compartments, whether by pipetting (*12–14*), sorting (*2, 8, 13, 15–17*), microfluidics-based deposition to microwells (*18*), or by dilution to emulsion-based droplets (*5, 19, 20*). As a consequence, the cost of preparing single cell RNA-seq libraries with these methods scales linearly with the numbers of cells processed. Although droplet-based methods are generally more cost-effective than well-based methods, they are nonetheless limited by linear cost-scaling, the need for specialized instrumentation, and an incompatibility with profiling nuclei or fixed cells (*21, 22*). Furthermore, droplet-based systems capture cells based on cell size, which may bias analyses of heterogeneous tissues.

We set out overcome these limitations with a new method for single cell RNA-seq based on the concept of combinatorial indexing. Combinatorial indexing uses split-pool barcoding to uniquely label a large number of single molecules or single cells, but without requiring the physical isolation of each molecule or cell. We previously used combinatorial indexing of single molecules (high molecular weight genomic DNA fragments) for both haplotype-resolved genome sequencing and *de novo* genome assembly (*23, 24*). More recently, we and others demonstrated <u>s</u>ingle cell <u>c</u>ombinatorial <u>i</u>ndexing (“sci”) to efficiently profile chromatin accessibility (sci-ATAC-seq) (*25*), genome sequences (sci-DNA-seq) (*26*), and genome-wide chromosome conformation (sci-Hi-C) (*27*) in large numbers of single nuclei.

Here we describe sci-RNA-seq, a straightforward method for profiling the transcriptomes of large numbers of single cells or nuclei per experiment, using off-the-shelf reagents and conventional instrumentation. For single cells, the protocol relies on methanol fixation, which can stabilize and preserve RNA in dissociated cells for weeks (*28*), thereby minimizing perturbations to cell state before or during processing. In this proof-of-concept, we apply sci-RNA-seq to profile the transcriptomes of ~16,000 mammalian cells in a single experiment, and show that we can separate synthetic mixtures of inter- or intraspecies cell types. We then apply sci-RNA-seq to *Caenorhabditis elegans* worms at the L2 stage, sequencing the transcriptomes of 42,035 single cells. This includes 37,734 somatic cells, effectively 50x coverage (*i.e.* oversampling) of this organism’s entire cellular content (762 cells at the L2 stage). From these data, we identify 27 distinct cell types, estimate their consensus expression profiles and define cell-type specific and selective genes, including for some fine-grained cell types that are present in only one or two cells per animal.

## Results

### Overview of method

In its current form, sci-RNA-seq relies on the following steps (**Fig. 1a**): (1) Cells are fixed and permeabilized with methanol (alternatively, cells are lysed and nuclei recovered), and then split across 96- or 384-well plates. (2) To introduce a first molecular index to the mRNA of cells within each well, we perform *in situ* reverse transcription (RT) with a barcode-bearing, well-specific polyT primer with unique molecular identifiers (UMI). (3) All cells are pooled and redistributed by fluorescence activated cell sorting (FACS) to 96- or 384-well plates in limiting numbers (*e.g.* 10-100 per well). (4) We then perform second strand synthesis, transposition with Tn5 transposase (Nextera), lysis, and PCR amplification. The PCR primers target the barcoded polyT primer on one end, and the Tn5 adaptor insertion on the other end, such that resulting PCR amplicons preferentially capture the 3’ ends of transcripts. Critically, these primers introduce a second barcode, specific to each well of the PCR plate. (5) Amplicons are pooled and subjected to massively parallel sequencing, resulting in 3’-tag digital gene expression profiles, with each read associated with two barcodes corresponding to the first and second rounds of cellular indexing (**Fig. 1b**).

**Figure 1:**
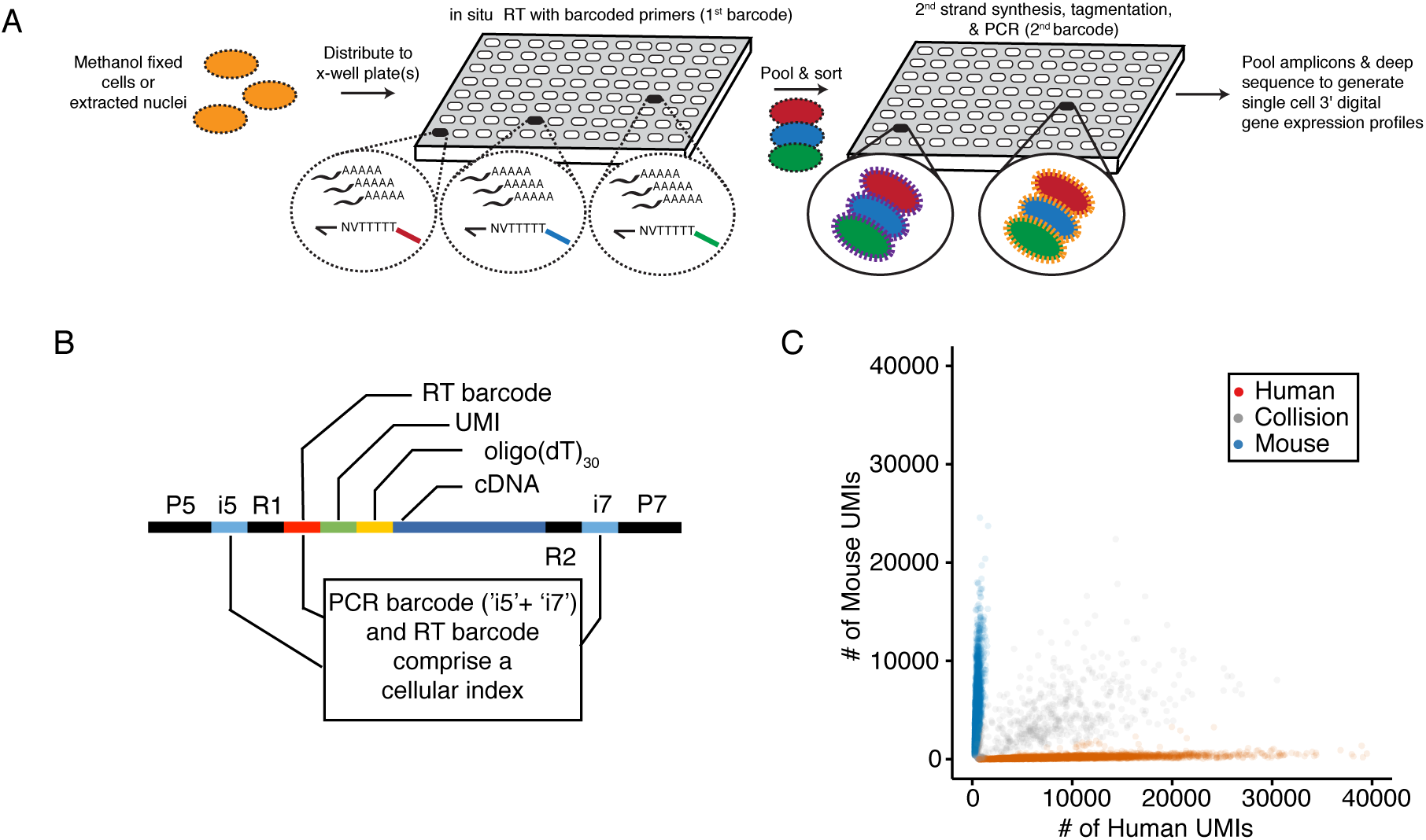
sci-RNAseq enables massively multiplexed single cell transcriptome profiling. a.) Schematic of the sci-RNA-seq workflow. Methanol-fixed cells or unfixed nuclei are split to one or more 96-well or 384-well plates for reverse transcription with different barcodes (first round of barcoding) in each well. Cells from different wells are pooled together and flow-sorted into one or more 96-well or 384-well plates for second strand synthesis, tagmentation and PCR with well-specific barcode combinations (second round of barcoding). The resulting PCR amplicons are pooled and deep sequenced to generate single cell 3’ digital gene expression profiles. b.) sci-RNA-seq library amplicons include Illumina adapters, PCR indices (i5 and i7), a reverse transcription barcode and UMI, in addition to the cDNA fragment to be sequenced. Read 1 covers the reverse transcription barcode (10 bp) and unique molecular identifier (UMI, 8 bp). Read 2 covers the cDNA fragment. The combination of the PCR indices and the reverse transcription barcode define a cellular index. c.) Scatter plot of unique human and mouse UMI counts from a 384 x 384 sci-RNA-seq experiment. This 384-well experiment included multiple different mixtures of cells (see Methods), but only cells originating from a well containing mixed human (HEK293T or HeLa S3) and mouse (NIH/3T3) cells during the first round of barcoding are plotted here. Inferred mouse cells are colored in blue; inferred human cells are colored in red, and “collisions” are colored in grey.

Because the overwhelming majority of cells pass through a unique combination of wells, each cell is effectively “compartmentalized” by the unique combination of barcodes that tag its transcripts. As we previously described, the rate of “collisions”, *i.e.* two or more cells receiving the same combination of barcodes, can be tuned by adjusting how many cells are distributed to the second set of wells (*25*). The sub-linear cost-scaling of combinatorial indexing follows from the fact that the number of possible barcode combinations is the product of the number of barcodes used at each stage. Consequently, increasing the number of barcodes used in the two rounds of indexing leads to an increased capacity for the number of cells that can be profiled and a lower effective cost per cell (**Fig. S1**). Additional rounds of molecular indexing can potentially offer even greater complexity and lower costs. Pertinent to experiments described below, we note that multiple samples (*e.g.* different cell populations, tissues, individuals, time-points, perturbations, replicates, etc.) can be concurrently processed within a single sci-RNA-seq experiment, simply by using different subsets of wells for each sample during the first round of indexing.

### Application of sci-RNA-seq to mammalian cells

To evaluate the scalability of this strategy, we performed a 384 well x 384 well sci-RNA-seq experiment. During the first round of indexing, half of the 384 wells contained pure populations of either human (HEK293T or HeLa S3) or mouse (NIH/3T3) cells, whereas half of the wells contained a mixture of human and mouse cells. After barcoded RT, cells were pooled and then sorted to a new 384 well plate for further processing, including the second round of barcoding and deep sequencing of pooled PCR amplicons. From this single experiment, we recovered 15,997 single cell transcriptomes.

At a read depth corresponding to ~25,000 reads per cell (~65% duplication rate), we recovered (on average) 11,024 UMIs per HEK293T cell, 7,832 UMIs per HeLa S3 cell, and 5,260 UMIs per NIH/3T3 cell. Cells originating in wells containing an interspecies mixture were readily assignable as human or mouse, with a rate of collision 16% (6.3% expected) (**Fig. 1c**). Species mixing experiments provide a means of estimating ‘impurities’ that might result from mRNA leakage between permeabilized cells. In this initial experiment, the rate of human reads appearing in mouse-assigned cells was 6.2%, and the rate of mouse reads appearing in human cells was 2.9%.

To reduce mRNA leakage as well as to increase the number of unique transcripts profiled per cell, we extensively optimized the protocol, most importantly the choice of reverse transcriptase and the washing steps. In an experiment that is representative of the culmination of these optimizations, we performed 96 well x 96 well sci-RNA-seq on five mammalian cell populations, split across 96 wells during the first round of barcoding: HEK293T cells (8 wells); HeLa S3 cells (8 wells); an intraspecies mixture of HEK293T and HeLa S3 cells (32 wells); and interspecies mixtures of HEK293T and NIH/3T3 cells (24 wells) or nuclei (24 wells). After barcoded reverse transcription across these 96 wells, cells were pooled, sorted to a new 96 well plate for a second round of barcoding by PCR, and then all amplicons pooled.

We deeply sequenced this library to a read depth corresponding to ~250,000 reads per cell (~88% duplication rate). We grouped reads that shared the same first and second round barcodes (inferring that each such group originated from the same single cell), which yielded 744 single cell and 175 single nucleus transcriptomes. Transcript tags were aligned to a combined human and mouse reference genome with STAR (*29*). Transcriptomes originating in ‘human only’ wells overwhelmingly mapped to the human genome (99%), including 51,311 UMIs per cell from HEK293T-only wells and 31,276 UMIs per cell from HeLa S3-only wells (on average), much higher transcript counts per cell than our original experiment. 81% of reads mapped to the expected strand of genic regions (47% exonic, 34% intronic), and the remainder to the unexpected strand of genic regions (9%) or to intergenic regions (10%). These proportions are similar to other studies (*17*). Whereas exonic reads show the expected enrichment at the 3’ ends of gene bodies, intronic reads do not, and may be the result of poly(dT) priming from poly(dA) tracts in heterogeneous nuclear RNA (**Fig. S2**). However, because like exonic reads they overwhelmingly derived from the expected strand of genic regions, we retained them in subsequent analyses.

In this experiment, transcriptomes originating in wells containing an interspecies mixture of human (HEK293T) and mouse (NIH/3T3) cells overwhelmingly mapped to the genome of one species or the other (289 of 294 cells), with only 5 clear ‘collisions’ (3.4% collision rate; 4.3% expected) (**Fig. 2a**). Excluding these collisions, we observed 24,454 UMIs per human cell and 17,665 UMIs per mouse cell (**Fig. 2b-c**), with an average of 1.9% and 3.3% of reads per cell mapping to the incorrect species, respectively, a level of impurity that is lower than our original experiment and comparable to the droplet-based Drop-Seq method (*19*).

**Figure 2:**
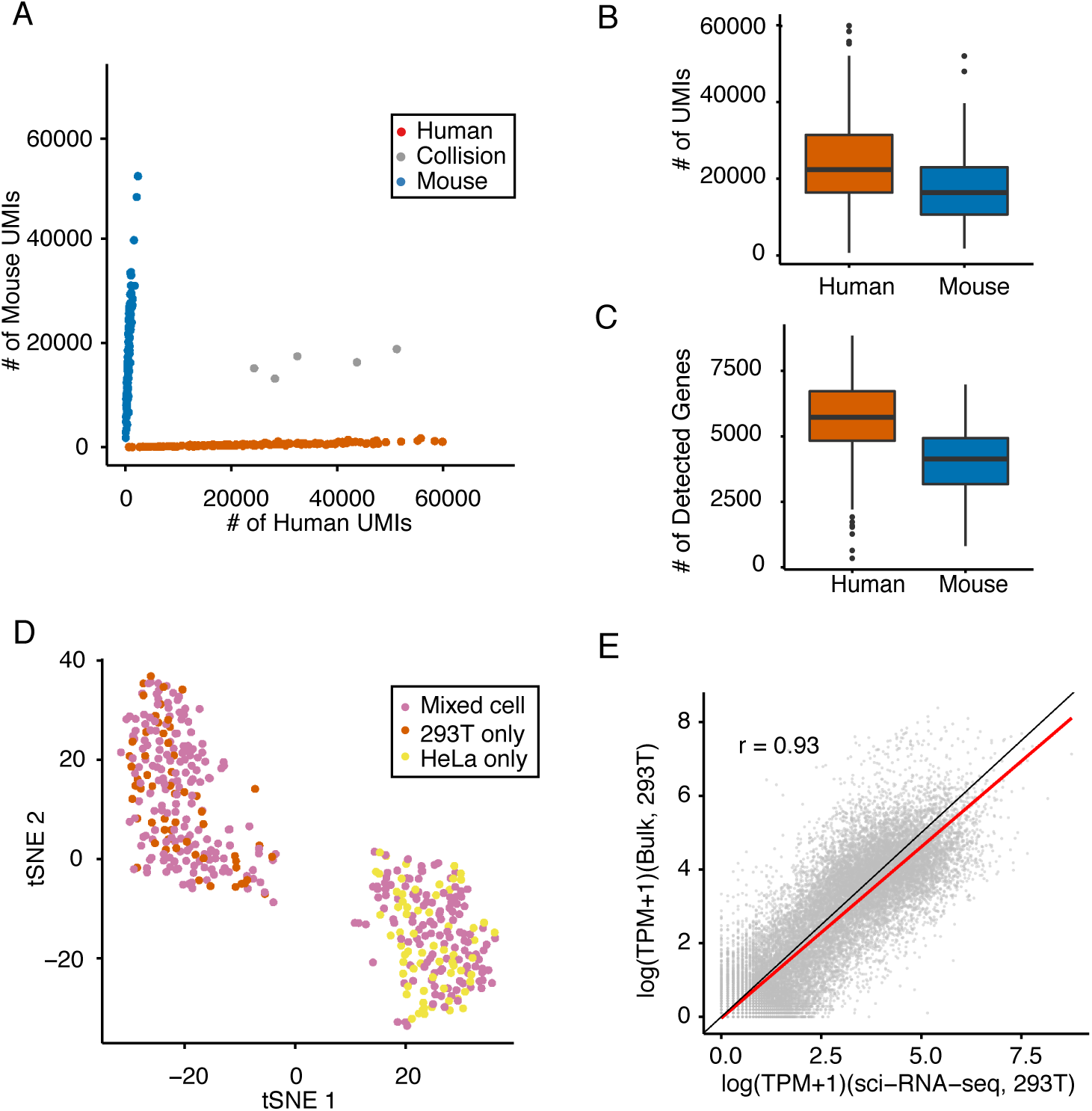
Representative results from an optimized protocol for sci-RNA-seq. a.) Scatter plot of unique human and mouse cell UMI counts from a 96-well sci-RNA-seq experiment. This 96-well experiment included multiple different mixtures of cells (see Methods), but only cells originating from a mixture of human (HEK293T) and mouse (NIH/3T3) are plotted here. Inferred mouse cells are colored in blue; inferred human cells are colored in red, and “collisions” are colored in grey. b.-c.) Boxplots showing the number of UMIs (b) and genes (c) detected per cell in interspecies mixing experiments. d.) tSNE plot of human cell line (HEK293T and HeLa S3) mixtures and pure populations. Cells originating in wells containing pure HEK293T (red), pure HeLa S3 (yellow) or a mixture of the two (pink) were all clustered together with tSNE. e.) Correlation between gene expression measurements from aggregated sci-RNA-seq data vs. bulk RNA-seq data, together with a linear regression line (red) and y=x line (black).

We next sought to confirm that we can separate cell types with sci-RNA-seq, focusing on the two human cell types (HEK293T and HeLa S3 cells) included in this experiment. We performed t-stochastic neighbor embedding (t-SNE) on standardized expression values of single cells derived from wells containing pure HEK293T cells, pure HeLa S3 cells, or mixed HEK293T and HeLa S3 cells. The mixed cells readily separate into two clusters, with one corresponding to HEK293T cells and the other to HeLa S3 cells (**Fig. 2d**). To further validate the result, we identified single nucleotide variants (SNVs) that distinguish HEK293T and HeLa S3 cells and found that our assignments are in agreement with the SNV-based assignments (**Fig. S3a**). Of note, the intronic reads alone are sufficient to separate HEK293T and HeLa S3 cells (**Fig. S3b**).

To evaluate whether sci-RNA-seq is biased compared to bulk measurements, we compared *in silico* aggregated transcriptomes from all 220 identified HEK293T cells against the output of a related bulk RNA-seq workflow (Tn5-RNA-seq (*30*)) without methanol fixation. The resulting estimates of gene expression were highly correlated (Pearson: 0.93; **Fig. 2e**).

### Application of sci-RNA-seq to mammalian nuclei

Protocols for dissociating tissues to single cell suspensions are labor intensive, do not easily standardize, potentially impact cell states, and potentially bias cell type composition. To address this, several groups have developed single nucleus RNA-seq protocols (since single nuclei are much more readily obtained than single cells), but these ‘one-nucleus-per-well’ approaches (*8, 10, 17*) do not efficiently scale.

We therefore evaluated sci-RNA-seq for profiling the transcriptomes of single nuclei extracted from a synthetic mixture of mouse (NIH/3T3) and human (HEK293T) cells. From the above-described experiment in which these nuclei were sorted to 24 of the 96 wells during the first round of barcoding, we recovered 175 single nucleus transcriptome profiles. Transcript tags were aligned to a combined human and mouse reference genome with STAR (*29*). As with single cells, reads associated with single nuclei mapped overwhelmingly to the genome of one species or the other (48 human nuclei; 124 mouse nuclei) with 3.4% collisions rate (4.3% expected) (**Fig. 3a**).

**Figure 3:**
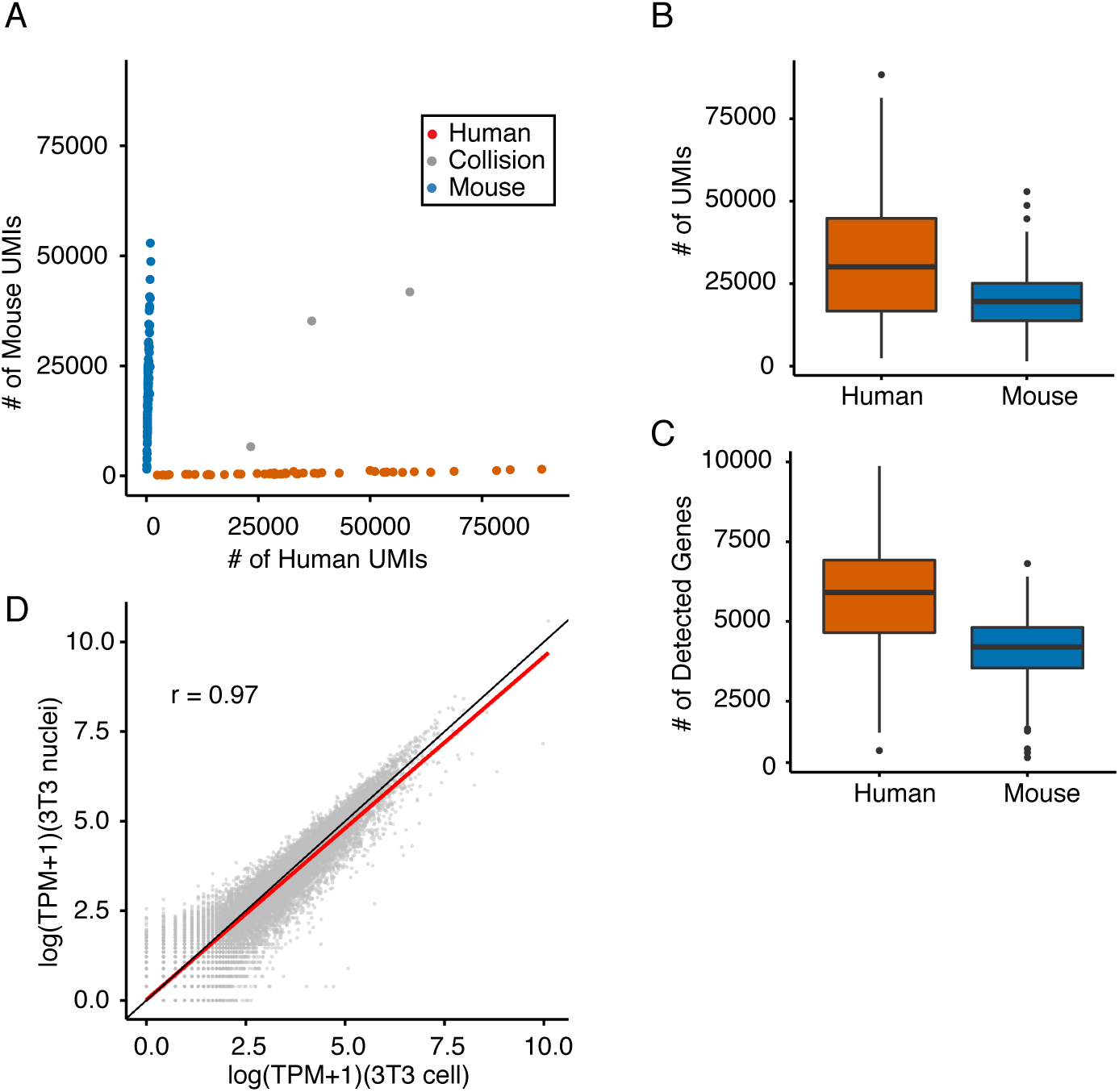
sci-RNA-seq is compatible with isolated nuclei as starting material. a.) Scatter plot of unique human and mouse nuclei UMI counts from a 96-well sci-RNA-seq experiment. This 96-well experiment included multiple different mixtures of cells (see Methods), but only cells originating from a mixture of human (HEK293T) and mouse (NIH/3T3) nuclei are plotted here. Inferred murine cells are colored in blue; inferred human cells are colored in red, and “collisions” are colored in grey. b.-c.) Boxplots showing the number of UMIs (b) and genes (c) detected per cell in nuclear sci-RNA-seq experiments. d.) Correlation between gene expression measurements in aggregated sci-RNA-seq profiles of NIH/3T3 cells vs. NIH/3T3 nuclei, together with a linear regression line (red) and y=x line (black).

Excluding collisions and at a read depth corresponding to ~210,000 reads per nucleus (~88% duplication rate), we observed 32,951 UMIs per human nucleus and 20,123 UMIs per mouse nucleus (**Fig. 3b-c**), with an average of 2.2% and 1.9% of reads per cell mapping to the incorrect species. 84% of reads mapped to the expected strand of genic regions (35% exonic, 49% intronic) and 16% to intergenic regions or to the unexpected strand of genic regions. These proportions are similar to previous descriptions of single nucleus RNA-seq (*17*), which together with the number of transcript molecules recovered indicate that sci-RNA-seq can flexibly and scalably profile the transcriptomes of either cells or nuclei. As a further check, we aggregated the transcriptomes of identified NIH/3T3 nuclei (*n* = 124) and compared them against the aggregated transcriptomes of identified NIH/3T3 cells (*n* = 129) and found the resulting estimates of gene expression to be highly correlated (Pearson: 0.97; **Fig. 3d**).

### Single cell RNA profiling of *C. elegans*

The roundworm *C. elegans* is the only multicellular organism for which all cells and cell types are defined, as is its entire developmental lineage (*31, 32*). As the extent to which contemporary single cell experimental and computational methods can comprehensively recover and distinguish cell types remains a matter of debate, we applied sci-RNA-seq to whole *C. elegans* larvae. Of note, the cells in *C. elegans* larvae are much smaller, more variably sized, and have markedly lower mRNA content than mammalian cells, and therefore represent a much more challenging test of sci-RNA-seq’s technical robustness.

We pooled ~150,000 larvae synchronized at the L2 stage and dissociated them into single-cell suspensions. We then performed *in situ* reverse transcription across 6 96-well plates (*i.e.* 576 first-round barcodes), with each well containing ~1,000 *C. elegans* cells and also 1,000 human cells (HEK293T) as internal controls. After pooling cells from all plates together, we sorted the mixture of *C. elegans* cells and HEK293T cells into 10 new 96-well plates for PCR barcoding (*i.e.* 960 second-round barcodes), gating on DNA content to distinguish between *C. elegans* and HEK293T cells. This sorting was carried out such that 96% of wells harbored only *C. elegans* cells (140 each), and 4% of wells harbored a mix of *C. elegans* and HEK293T cells (140 *C. elegans* cells and 10 HEK293T cells each).

This single experiment yielded 42,035 *C. elegans* single-cell transcriptomes (number of UMI counts for protein-coding transcripts > 100). 93.7% of reads mapped to the expected strand of genic regions (91.7% exonic, 2.0% intronic). At a sequencing depth of ~20,000 reads per cell and a duplication rate of 79.9%, we identified a median of 575 UMIs mapping to protein coding genes per *C. elegans* cell (mean of 1,121 UMIs per cell), likely reflective of the lower mRNA content of *C. elegans* cells relative to mammalian cells (**Fig. S4a**). Importantly, control wells containing both *C. elegans* and HEK293T cells demonstrated clear separation between the two species (**Fig. S4b**).

Semi-supervised clustering analysis segregated the cells into 29 distinct groups with the largest containing 13,205 cells (31.4%) and the smallest containing only 131 cells (0.3%) (**Fig. 4a**). Somatic cell types comprised 37,734 cells. We identified genes that were expressed specifically in a single cluster, and by comparing those genes to expression patterns reported in the literature assigned the clusters to cell types (**Table S2**). Of the 29 clusters, 21 represented exactly one literature-defined cell type, 5 contained >1 distinct cell types, and 3 contained cells of unclear type. Neurons, which were present in 7 clusters from the global analysis, were independently subclustered, revealing 10 major neuronal subtypes (**Fig. 4b**).

**Figure 4:**
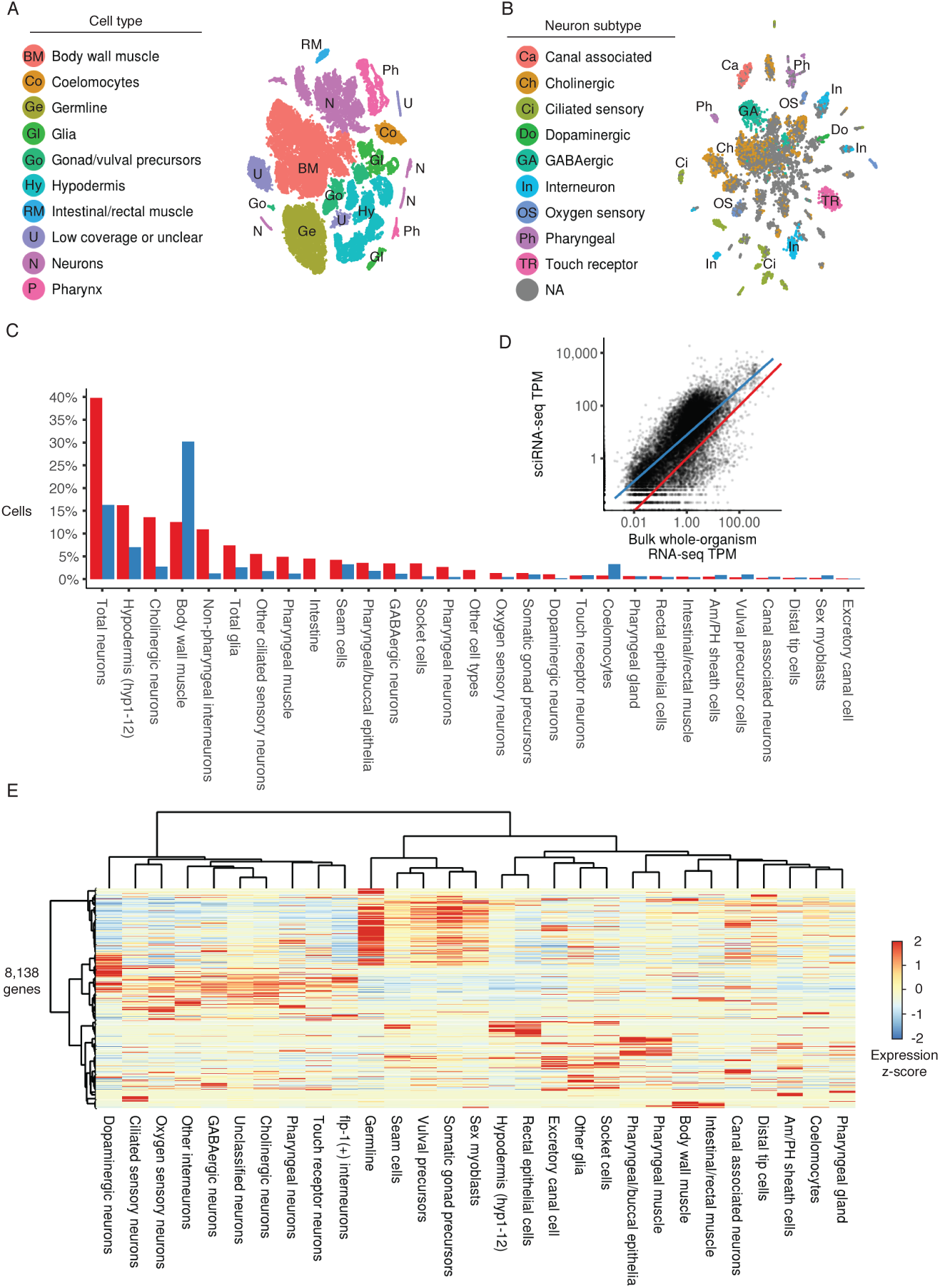
A single sciRNA-seq experiment provides a near-comprehensive view of the single cell transcriptomes comprising the *C. elegans* larva. a) t-SNE visualization of the high-level cell types identified. b) t-SNE visualization of neuronal subtypes. Dimensionality reduction and clustering was applied to cells in neuronal clusters shown in (a). c) Bar plot showing the proportion of somatic cells profiled with sci-RNA-seq that could be identified as belonging to each cell type compared to the proportion of cells from that type present in an L2 *C. elegans* individual. d) Scatter plot showing the log-scaled transcripts per million (TPM) of genes in the aggregation of all sci-RNA-seq reads (x axis) or in bulk RNA-seq (y axis; geometric mean of 3 experiments), e) Heatmap showing the relative expression of genes in consensus transcriptomes for each cell type estimated by sci-RNA-seq. Genes are included if they have a size-factor-normalized mean expression of  0.05 in at least one cell type (8,318 genes in total). The raw expression data (UMI count matrix) is log-transformed, column centered and scaled (using the R function scale), and the resulting values are clamped to the interval [−2, 2].

Overall, the global and neuron-specific clustering analyses allowed for the construction of expression profiles of 27 well-defined cell types (**Fig 4e**; not counting “unclassified neurons” and “other glia”). 11 of these cell types are represented by 1 to 6 cells in an individual L2 worm. Examples of these fine-grained cell types include the distal tip cells of the developing gonad (2 cells), the excretory canal cell (1 single cell) and canal associated neurons (2 cells), and the amphid/phasmid sheath cells (4 glial cells).

Each *C. elegans* animal contains exactly the same number of cells, which are generated deterministically in a defined lineage (*31, 32*). Comparing the observed proportions of each cell type to their known frequencies in L2 larvae showed that sci-RNA-Seq captured many cell types at or near expected frequencies (**Fig. 4c**; 15 types had abundance ≥ 50% of the expectation). Body wall muscle was 2.4-fold more abundant in our data than in the animal, probably reflecting its ease of dissociation. Only one abundant cell type, the intestinal cells, was absent. We speculate that this was due to our gating strategy during FACS, which excluded cells based on DAPI signal. Intestinal cells at the L2 stage are polyploid, containing 8 copies of each (haploid) chromosome.

Previous analyses of single-cell transcriptomes have shown that cells can often be distinguished with relatively light sequencing per cell on the basis of a small set of highly specific genes (*33*). This raised the possibility that despite being able to detect many distinct cell types in the worm, our molecular definition for each would be incomplete. However, we observed that half of all *C. elegans* protein-coding genes were expressed in at least 80 cells in the full dataset, and 64% of protein-coding genes were expressed in at least 20 cells. As many genes are expressed specifically in embryos or adults, our gene expression measurements likely include most of the protein-coding genes that are truly expressed in L2 *C. elegans*. The “whole-worm” expression profile derived by aggregating all sci-RNA-Seq reads correlated well with published whole-organism bulk RNA-seq (*34*) for L2 *C. elegans* (**Fig. 4d,** Spearman ρ = 0.795). Furthermore, consensus expression profiles for each cell type segregated as expected in a hierarchical clustering analysis (**Fig. 4e**). Thus, despite the fact that sci-RNA-Seq captures a minority of transcripts in each cell, our ‘oversampling’ of the cellular composition of the organism (*i.e.* 37,734 somatic cells, effectively 50x coverage of the L2 worm’s 762 somatic cells), enables us to construct faithful expression profiles for individual cell types.

## Discussion

We describe a new method for single cell RNA-seq based on the principle of combinatorial indexing of cells or nuclei. At the scale described here (*e.g.* 576 x 960 indexing), sci-RNA-seq can be applied to profile the transcriptomes of tens-of-thousands of single cells per experiment. Library preparation can be completed by a single individual in two days. The cost of library preparation, currently $0.03-$0.20 per cell, is dominated by enzymes. The method relies on off-the-shelf reagents and widely available instrumentation (*e.g.* FACS, sequencer).

sci-RNA-seq has several practical advantages over contemporary alternatives:

First, it is compatible (and indeed relies on) cell fixation, which can minimize perturbations to cell state or RNA integrity before or during processing.

Second, sci-RNA-seq facilitates the concurrent processing of multiple samples (*e.g.* corresponding to different cell populations, tissues, individuals, time-points, perturbations, replicates, etc.) within a single experiment, simply by using different subsets of wells for each sample during the first round of indexing. This may have the benefit of reducing batch effects, relative to platforms requiring the serial processing of samples, an area of paramount concern for the single cell RNA-seq field (*35*). Furthermore, given that the second barcode is introduced after flow sorting, it is also possible to associate wells on the PCR plate with FACS-defined subpopulations (*e.g.* corresponding to cell size, cell cycle, immunostaining, etc.), as we did while sorting mixed HEK293T and *C. elegans* cells.

Third, as we show, sci-RNA-seq is readily compatible with the processing of nuclei. Single nucleus RNA-seq is possible with “well-per-well” methods but is not currently supported on droplet-based platforms. The ability to process nuclei, rather than cells, may be particularly important for tissues for which unbiased cell disaggregation protocols are not well established (which may be most tissues).

Fourth, sci-RNA-seq is highly scalable. Here we demonstrate up to 576 × 960 combinatorial indexing, which enables the generation of ~5 × 10^4^ single cell transcriptomes. While beyond the scope of this proof-of-concept, one can imagine simply using more barcoded RT and PCR primers (*e.g.* 1,536 × 1,536 combinatorial indexing), which would enable processing of many more cells with sub-linear scaling of the cost per cell. A complementary approach is simply to introduce additional rounds of indexing, *e.g.* by using indexed Tn5 complexes during *in situ* transposition (*24, 25*). With 384 × 384 × 384 combinatorial indexing, we can potentially uniquely barcode the transcriptomes of >10 million cells within a single experiment.

In this proof-of-concept, we apply sci-RNA-Seq to generate the first catalog of single cell transcriptomes at the scale of a whole organism. In a single experiment, we generated ~50X coverage of the somatic cellular composition of L2 *C. elegans*, detecting 27 cell types and constructing consensus transcriptional profiles for each. While not all cell types are detected at expected frequencies, 15 cell types had abundance ≥ 50% of the expectation. Half of *C. elegans* protein-coding genes were expressed in at least 80 cells in the full dataset, demonstrating that our expression profile for this stage of *C. elegans* development is mostly comprehensive. Taken together, these analyses show that sci-RNA-Seq constitutes a reliable platform for molecular dissection of complex tissues and potentially whole organisms.

sci-RNA-seq further expands the repertoire of single cell molecular phenotypes that can be resolved by combinatorial indexing (which now includes mRNA, chromatin accessibility, genome sequence, and chromosome conformation). Looking forward, we anticipate that additional forms of single cell profiling can be achieved with combinatorial indexing. Provided that multiple aspects of cellular biology can be concurrently barcoded, combinatorial indexing may also facilitate the scalable generation of ‘joint’ single cell molecular profiles (*e.g.* RNA-seq and ATAC-seq from each of many single cells).

## Materials & Methods

### Cell Culture

All cells were cultured at 37°C with 5% CO_2_, and were maintained in high glucose DMEM (Gibco cat. no. 11965) supplemented with 10% FBS and 1X Pen/Strep (Gibco cat. no. 15140122; 100U/ml penicillin, 100μg/ml streptomycin). Cells were trypsinized with 0.25% typsin-EDTA (Gibco cat. no. 25200-056) and split 1:10 three times a week.

### Generation of whole *C. elegans* cell suspensions

A *C. elegans* strain (RW12139 *stIs11435(unc-120::H1-Wcherry;unc-119(+));unc-119(tm4063)*) carrying an integrated Punc-120::mCherry gene in a wild type background was used in all experiments. A synchronized L2 population was obtained by two cycles of bleaching gravid adults to isolate fertilized eggs allowing the eggs to hatch in the absence of food to generate a population of starved L1 animals. Around 150,000 L1 larvae were plated on each 100 mm petri plate seeded with NA22 bacteria and incubated at 24 °C for 15 hr to produce early L2 larvae. Dissociated cells were recovered following a published protocol (*36*) with modification. Specifically, L2 stage worms were collected by adding 10 ml sterile ddH2O to each plate. The collected L2s were pelleted by centrifugation at 1300 g for 1 min. The larval pellet was washed five times with sterile ddH2O to remove bacteria. The resulting pellet was transferred to a 1.6 ml microcentrifuge tube. Around 40 μl of the final compact pellet was used for each cell dissociation experiment. The worm pellet was treated with 250 μl of SDS-DTT solution (20 nM HEPES pH8, 0.25% SDS, 200 mM DTT, 3% sucrose) for 4 min. Immediately after SDS-DTT treatment, egg buffer was added to the SDS-DTT treated worms. Worms were pelleted at 500 g for 1 min, then washed 5 times with Egg buffer (118 mM NaCl. 48 mM KCl. 3 mM CaCl2. 3 mM MgCl2. 5 mM HEPES (pH 7.2)). Pelleted SDS-DTT treated worms were digested with 200 μl of 15 mg/ml pronase (Sigma-Aldrich, St. Louis, MO) for 20 min. The treated worms were broken up to release cells by pipetting up and down with 21G1 1/4 needle. When sufficient single cells were observed the reaction was stopped by adding 900 μl L-15 medium containing 10% fetal bovine serum. Cells were separated from worm debris by centrifuging the pronase-treated worms at 150 g for 5 min at 4°C. The supernatant was transferred to 1.6 ml microcentrifuge tube and centrifuged at 500 g for 5 min at 4°C. The cell pellet was washed twice with egg-buffer containing 1% BSA.

### Sample Processing

All cell lines were trypsinized, spun down at 300x**g** for 5 min (4°C). and washed once in 1X PBS. *C. elegans* cells were dissociated as described above.

For sci-RNA-seq on whole cells, 5M cells were fixed in 5 mL ice-cold 100% methanol at −20 °C for 10 min, washed twice with 1 ml ice-cold 1X PBS containing 1% Diethyl pyrocarbonate (0.1% for *C. elegans* cells) (DEPC; Sigma-Aldrich), washed three times with 1 mL ice-cold PBS containing 1% SUPERase In RNase Inhibitor (20 U/μL, Ambion) and 1% BSA (20 mg/ml, NEB). Cells were resuspended in wash buffer at a final concentration of 5000 cells/ul. For all washes, cells were pelleted through centrifugation at 300x**g** for 3 min, at 4°C.

For sci-RNA-seq on nuclei, 5M cells were combined and lysed using 1 mL ice-cold lysis buffer (10 mM Tris-HCl, pH 7.4, 10 mM NaCl, 3 mM MgCl2 and 0.1% IGEPAL CA-630 from (*37*)), modified to also include 1% SUPERase In and 1% BSA). The isolated nuclei were then pelleted, washed twice with 1 mL ice-cold 1X PBS containing 1% DEPC, twice with 500 μL cold lysis buffer, once with 500 μL cold lysis buffer without IGEPAL CA-630, and then resuspended in lysis buffer without IGEPAL CA-630 at a final concentration of 5000 nuclei/μL. For all washes, nuclei were pelleted through centrifugation at 300x**g** for 3 min. at 4°C).

For cell-mixing experiments, trypsinized cells were counted and the appropriate number of cells from each cell line were combined prior to fixation or lysis. Fixed cells or nuclei were then distributed into 96- or 384-well plates (see **Table S1)**. For each well, 1,000-10,000 cells or nuclei (2 μL) were mixed with 1 μl of 25 μM anchored oligo-dT primer (5′-ACGACGCTCTTCCGATCTNNNNNNNN[10bp index]TTTTTTTTTTTTTTTTTTTTTTTTTTTTTTVN −3′, where “N” is any base and “V” is either “A”, “C” or “G”; IDT) and 0.25 μL 10 mM dNTP mix (Thermo), denatured at 55°C for 5 min and immediately placed on ice. 1.75 μL of first-strand reaction mix, containing 1 μL 5X Superscript IV First-Strand Buffer (Invitrogen), 0.25 μl 100 mM DTT (Invitrogen), 0.25 μl SuperScript IV reverse transcriptase (200 U/μl, Invitrogen), 0.25 μL RNaseOUT Recombinant Ribonuclease Inhibitor (Invitrogen), was then added to each well. Reverse transcription was carried out by incubating plates at 55°C for 10 min, and was stopped by adding 5 μl 2X stop solution (40 mM EDTA, 1 mM spermidine) to each well. All cells (or nuclei) were then pooled, stained with 4',6-diamidino-2-phenylindole (DAPI, Invitrogen) at a final concentration of 3 μM, and sorted at varying numbers of cells/nuclei per well (depending on experiment; see **Table S1**) into 5 uL buffer EB using a FACSAria III cell sorter (BD). 0.5 μl mRNA Second Strand Synthesis buffer (NEB) and 0.25 μl mRNA Second Strand Synthesis enzyme (NEB) were then added to each well, and second strand synthesis was carried out at 16°C for 150 min. The reaction was then terminated by incubation at 75°C for 20 min.

Tagmentation was carried out on double-stranded cDNA using the Nextera DNA Sample Preparation kit (Illumina). Each well was mixed with 5 ng Human Genomic DNA (Promega), as carrier to avoid over-tagmentation and reduce losses during purification, 5 μL Nextera TD buffer (Illumina) and 0.5 μL TDE1 enyzme (Illumina), and then incubated at 55 °C for 5 min to carry out tagmentation. Note that because the PCR primers used to amplify libraries are specific to the RT products, tagmented carrier genomic DNA are not appreciably amplified or sequenced. The reaction was then stopped by adding 12 μL DNA binding buffer (Zymo) and incubating at room temperature for 5 min. Each well was then purified using 36 uL AMPure XP beads (Beckman Coulter), eluted in 16 μL of buffer EB (Qiagen), then transferred to a fresh multi-well plate.

For PCR reactions, each well was mixed with 2μL of 10 μM P5 primer (5′-AATGATACGGCGACCACCGAGATCTACAC[i5]ACACTCTTTCCCTACACGACGCTCTTCCGATCT-3′), 2 μL of 10 μM P7 primer (5′-CAAGCAGAAGACGGCATACGAGAT[i7]GTCTCGTGGGCTCGG-3′), and 20 μL NEBNext High-Fidelity 2X PCR Master Mix (NEB). Amplification was carried out using the following program: 75°C for 3 min, 98°C for 30 sec, 18-22 cycles of (98°C for 10 sec, 66°C for 30 sec, 72°C for 1 min) and a final 72°C for 5 min. After PCR, samples were pooled and purified using 0.8 volumes of AMPure XP beads. Library concentrations were determined by Qubit (Invitrogen) and the libraries were visualized by electrophoresis on a 6% TBE-PAGE gel. Libraries were sequenced on the NextSeq 500 platform (Illumina) using a V2 75 cycle kit (Read 1: 18 cycles, Read 2: 52 cycles, Index 1: 10 cycles, Index 2: 10 cycles).

### Read alignments and construction of gene expression matrix

Base calls were converted to fastq format and demultiplexed using Illumina’s bcl2fastq/ 2.16.0.10 tolerating one mismatched base in barcodes (edit distance (ED) < 2). Demultiplexed reads were then adaptor clipped using trim_galore/0.4.1 with default settings. Trimmed reads were mapped to the human reference genome (hg19), mouse reference genome (mm10), *C.elegans* reference genome (PRJNA13758) or a chimeric reference genome of hg19, mm10 and PRJNA13758, using STAR/v 2.5.2b (*38*) with default settings and gene annotations (GENCODE V19 for human; GENCODE VM11 for mouse, WormBase PRJNA13758.WS253.canonical_gene set for *C.elegans*). Uniquely mapping reads were extracted, and duplicates were removed using the unique molecular identifier (UMI) sequence (ED < 2, including insertions and deletions), reverse transcription (RT) index, and read 2 end-coordinate (*i.e.* reads with identical UMI, RT index, and tagmentation site were considered duplicates). Finally, mapped reads were split into constituent cellular indices by further demultiplexing reads using the RT index (ED < 2, including insertions and deletions). For mixed-species experiment, the percentage of uniquely mapping reads for genomes of each species was calculated. Cells with over 85% of UMIs assigned to one species were regarded as species-specific cells, with the remaining cells classified as mixed cells. The collision rate was calculated as twice the ratio of mixed cells (as we are blind to any collisions involving cells of the same species). For gene body coverage analysis of exonic reads, the split human and mouse single cell SAM files were concatenated and exonic reads were selected and analyzed using RSEQC/2.6.1, using BED annotation files downloaded from the UCSC Golden Path. For read position analysis for intronic reads, the split human and mouse single cell SAM files were concatenated and intronic reads were selected; the fractional position of each intronic read along the genomic distance between the TSS and transcript terminus was calculated, and these values used to generate a density plot.

To generate digital expression matrices, we calculated the number of strand-specific UMIs mapping to the exonic and intronic regions of each gene, for each cell; generally, fewer than 3% of total UMIs strand-specifically mapped to multiple genes. For multi-mapped reads, reads were assigned to the closest gene, except in cases where another intersected gene fell within 100 bp to the end of the closest gene, in which case the read was discarded. For most analyses we included both intronic and exonic UMIs in per-gene single-cell expression matrices.

### t-SNE visualization of HEK293T cells and HeLa S3 cells

We visualized the clustering of sci-RNA-seq data from populations of pure HEK293T, pure HeLa S3 and mixed HEK293T + HeLa S3 cells using t-Distributed Stochastic Neighbor Embedding (tSNE). The top 3,000 genes with the highest variance in the digital gene expression matrix for these cells were first given as input to Principal Components Analysis (PCA). The top 10 principal components were then used as the input to t-SNE, resulting in the two-dimensional embedding of the data show in **Fig. 2D**. The process was repeated using only intronic reads (**Fig. S3B**). For this analysis, the top 2,000 (instead of 3,000) highly variable genes were used as input to PCA; all other parameters remained unchanged.

### Genotyping of single HeLa cells by 3’ tag sequences

HeLa S3 cell identity was verified on the basis of homozygous alleles not present in the hg19 assembly, using a callset derived from (39). Single-cell BAM files (with cellular indices encoded in the “read_id” field) were concatenated, and then processed as follows using a python wrapper of the samtools API (i.e. pysam). For each homozygous alternate SNV overlapping with a GENCODE V19 defined gene (n = 865,417) in the HeLa S3 variant callset, we computed the fraction of matching (i.e. HeLa S3 specific) alleles, and computed this value for all cells where at least 1 read containing a polymorphic site. We then re-plotted in R the tSNE visualization shown in **
Fig. 2D**, now colored by the relative fraction of homozygous alternate alleles called for each cell.

### Analysis of *C. elegans* whole-organism sci-RNA-seq experiment

A digital gene expression matrix was constructed from the raw sequencing data as described above. The dimensionality of this matrix was reduced first with PCA (40 components) and then with t-SNE, giving a two-dimensional representation of the data. Similar to the approach in (*40*), cells in this two-dimensional representation were clustered using the density peak algorithm (*41*) as implemented in Monocle 2. Genes specific to each cluster were identified and compared to microscopy-based expression profiles reported in the literature (**Table S2**), allowing the distinct cell types represented in each cluster to be identified. Based on these results, we manually merged two clusters that both corresponded to body wall muscle, and manually split two clusters that included hypodermis, somatic gonad cells, and glia. Seven clusters exclusively contained neurons. We identified neuronal subtypes applying PCA, t-SNE, and density peak clustering to this subset of cells using the same approach as for the global cluster analysis.

Consensus expression profiles for each cell type were constructed by first dividing each column in the gene-by-cell digital gene expression matrix by the cell's size factor and then for each cell type, taking the mean of the normalized UMI counts for the subset of cells assigned to that cell type. These mean normalized UMI counts were then re-scaled to transcripts per million. Cells that were part of a cluster corresponding to a cell type but did not express any of the marker genes used to define that cell type were excluded when generating these consensus expression profiles.

### Comparing sci-RNA-seq and bulk RNA-seq data for HEK293T cells

To compare aggregated sci-RNA-seq single cell transcriptomes with bulk RNA-seq, we performed bulk RNA-seq using a modified protocol (*30*). In brief, 500 ng total RNA extracted from three biological replicate HEK293T samples (extraction using RNeasy kit (Qiagen)) with the RNeasy kit (Qiagen) were used for reverse transcription following the standard SuperScript II protocol. 500 ng total RNA (in 9 μL water) was mixed with 2 μL 25 uM oligo-dT(VN) (5′-ACGACGCTCTTCCGATCTNNNNNNNN[10bp index]TTTTTTTTTTTTTTTTTTTTTTTTTTTTTTVN −3′, where “N” is any base and “V” is either “A”, “C” or “G”; IDT) and 1 μL 10 mM dNTPs, then incubated at 65°C for 5 min Following incubation, 8 μL reaction mix (4 μL 5X Superscript II First-Strand Buffer, 2 μl 100 mM DTT, 1 μl SuperScript II reverse transcriptase, 1 μL RnaseOUT) was added. Reactions were incubated at 42°C for 50 min and terminated at 70°C for 15 min. For second strand synthesis, 2 μL RT product was mixed with 6.5 μL water, 1 μL mRNA Second Strand Synthesis buffer (NEB) and 0.25 μl mRNA Second Strand Synthesis enzyme (NEB). Second strand synthesis was carried out at 16°C for 150 min, followed by 75°C for 20 min. Tagmentation was carried out by adding 10 μL Nextera TD buffer, 1 μL Nextera Tn5 enyzme and incubating at 55°C for 5 min. Tagmented cDNA was purified using a Clean & Concentrator^TM^-100 kit (Zymo) and eluted in 16 uL buffer EB. PCR, purification, and quantification were then performed as detailed above.

For comparing single cell RNA-seq and bulk RNA-seq, single cell gene counts of exonic reads and intronic reads were added for the same gene from sci-RNA-seq of pure HEK293T cells as well as HEK293T cells identified from HEK293T and NIH/3T3 mixed cells. Counts for bulk RNA-seq of HEK293T cells were extracted based on the RT barcode and aggregated separately, again adding exonic and intronic read counts per gene. Transcript counts were converted to transcripts per million (TPM) and then transformed to log(TPM + 1). Pearson correlation coefficients were calculated between the aggregated sci-RNA-seq and bulk RNA-seq data using R.

### Cost estimation

For 576 × 960 sci-RNA-seq, reagent costs are largely enzyme-driven and include SuperScript IV reverse transcriptase ($934), second strand synthesis mix ($750), Nextera Tn5 enzyme ($5,000), NEBnext master mix ($1,150) and other reagents and plates ($500). If we sort 60 cells per well (assuming recovery rate is 100%) for 960 wells (5% collision rate), then the reagent cost of library preparation per single cell is around $0.14 (expected yield of around 55,000 cells). However, it is worth noting that simply increasing the number of cells sorted per well also decreases costs (*e.g.* sort 150 cells to each well would yield around 140,000 cells at a cost of $0.05 per cell), but also results in an increased collision rate (12%). Alternatively, by increasing to 1,536 barcodes during the first (RT-based) round of indexing, we can sort up to 320 cells per well at a 10% collision rate, thereby reducing the cost per cell to less than $0.025 per cell. Straightforward reductions in reaction volumes at all steps may also lead to further reductions in costs, as would additional rounds of molecular indexing.

## Acknowledgements

We thank members of the Shendure, Trapnell and Waterston labs for helpful discussions; D. Prunkard, and L. Gitari in the Pathology Flow Cytometry Core Facility for their exceptional assistance in flow sorting; J. Rose, D. Maly, L. VandenBosch, and T. Reh lab for sharing the NIH/3T3 cell line. HeLa S3 cells were used as part of this study. Henrietta Lacks, and the HeLa cell line that was established from her tumor cells in 1951, have made significant contributions to scientific progress and advances in human health. We are grateful to Henrietta Lacks, now deceased, and to her surviving family members for their contributions to biomedical research. This work was funded by grants from the NIH (DP1HG007811 and R01HG006283 to JS; U41HG007355 and R01GM072675 to RHW) and the W. M. Keck Foundation (to CT and JS). DAC was supported in part by T32HL007828 from the National Heart, Lung, and Blood Institute. JS is an Investigator of the Howard Hughes Medical Institute.

---

**Figure S1.**
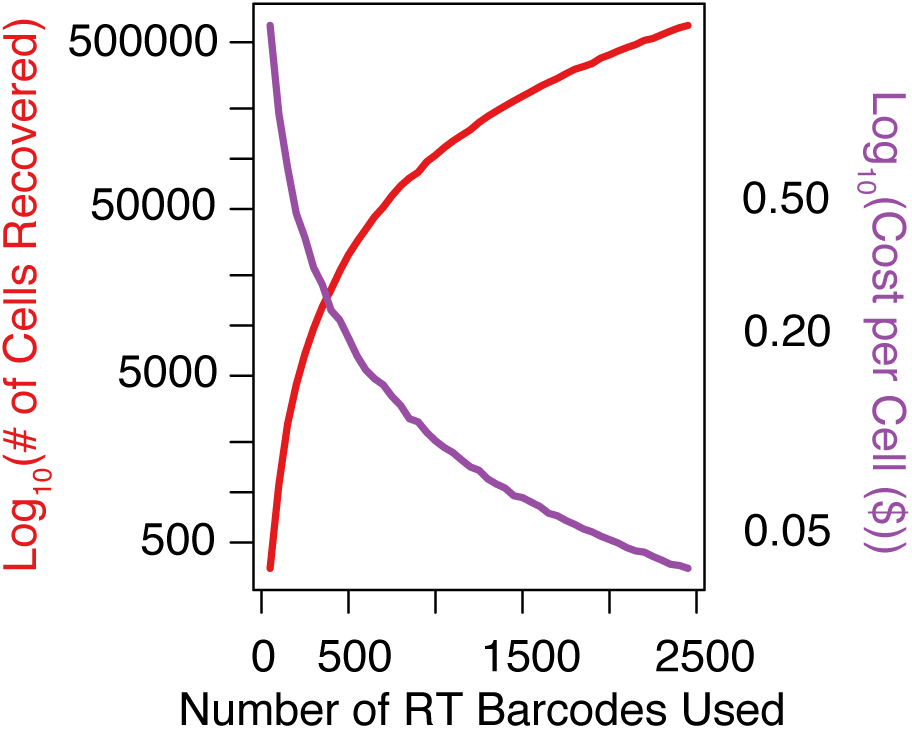
Combinatorial indexing with increasing numbers of reverse-transcription barcodes enables sublinear scaling of cost per cell. Plot showing how detection capacity (i.e. the number of cells detected in a sci-RNA-seq experiment, red) and cost per cell (blue) vary as a function of the number of cellular indices used, assuming a collision rate of 5%.

**Figure S2.**
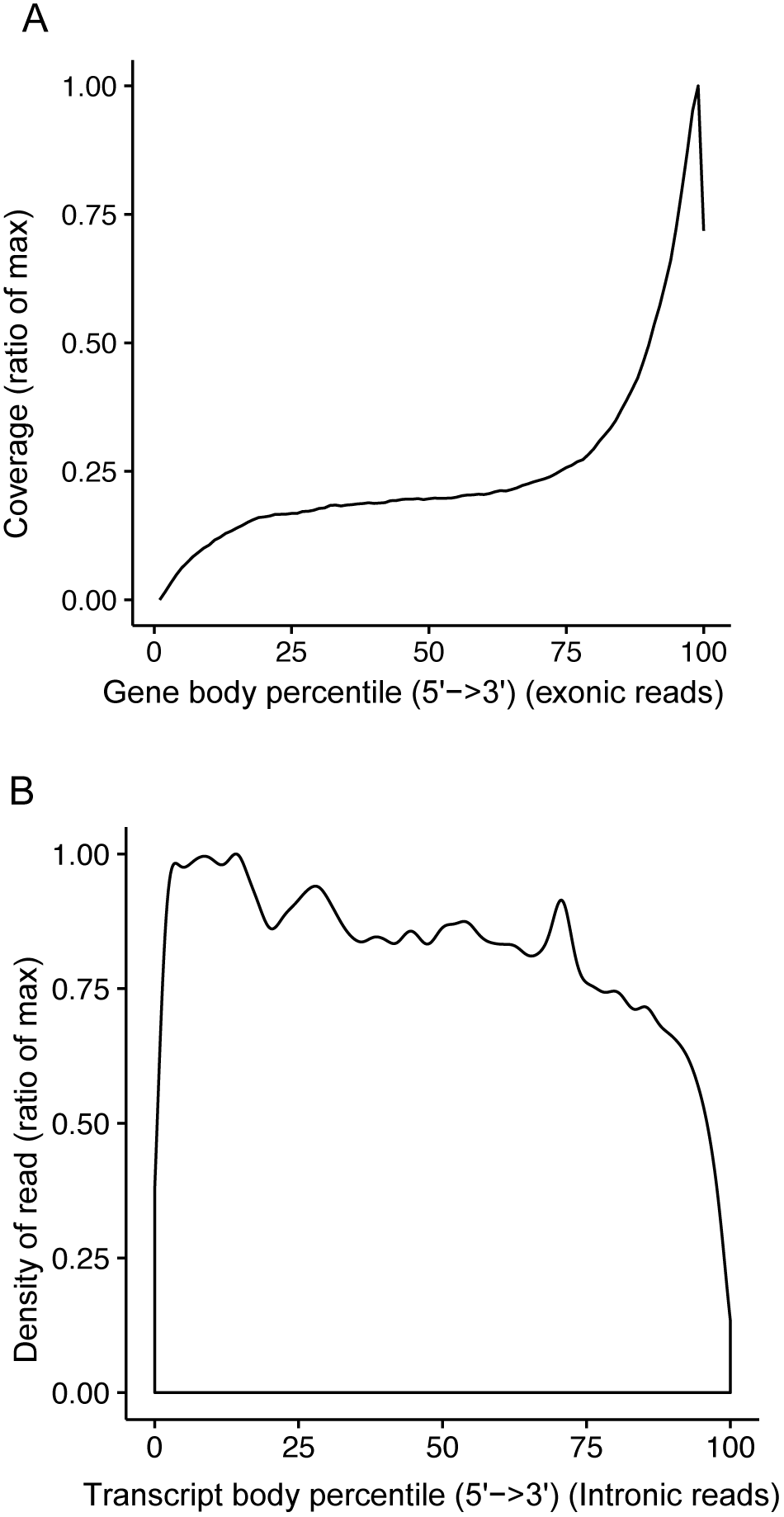
Metagene plot for unique intronic and exonic sci-RNA-seq reads. A.) Sci-RNA-seq on fixed cells demonstrates 3’-biased exonic coverage along gene body (intronic region exluded). B.) Density plot for the intronic reads number mapping to different percentiles of transcript body (intronic region included). y-axis is scaled to the ratio of max.

**Figure S3.**
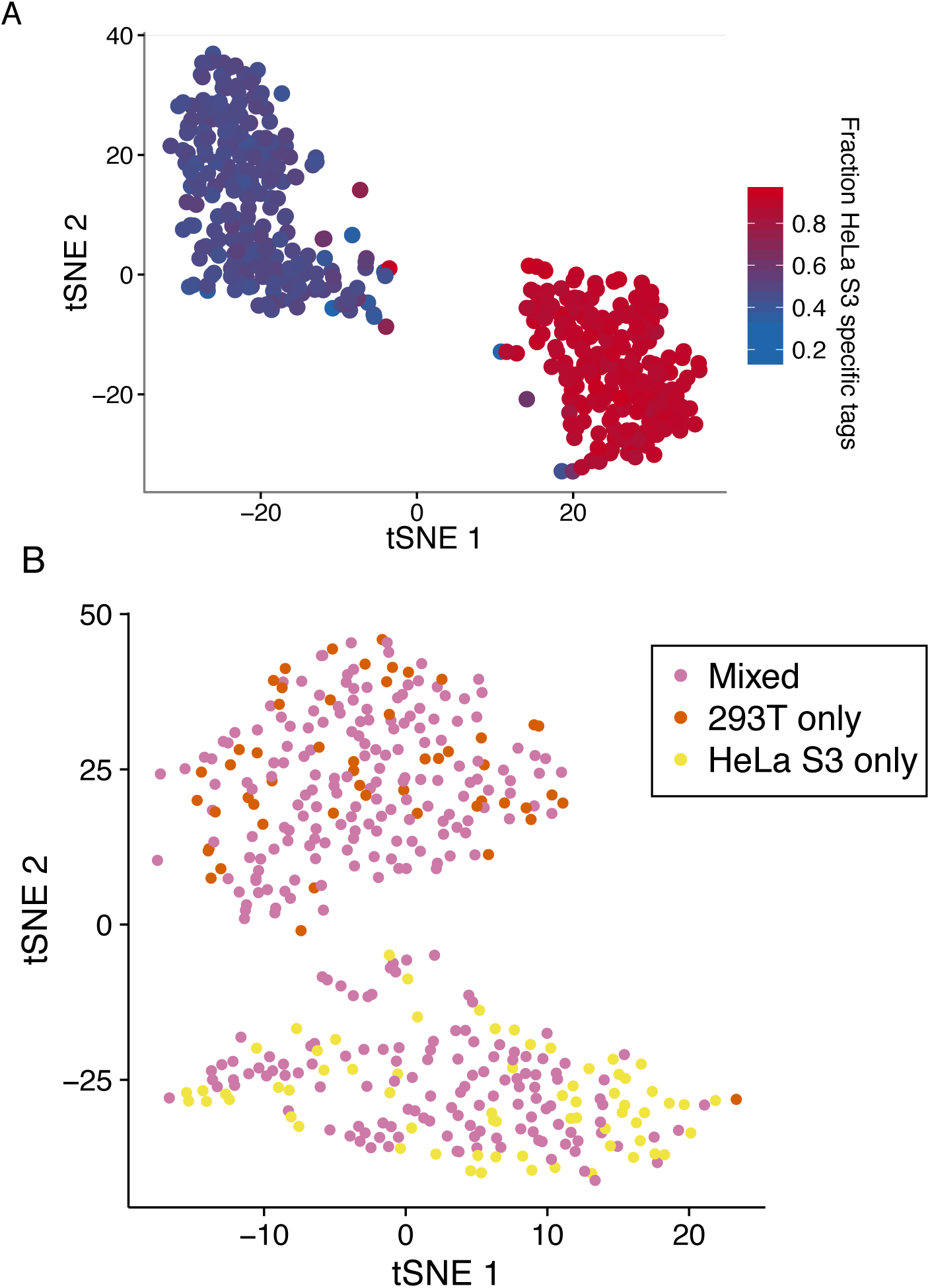
Quality control for sci-RNA-seq experiments using synthetic mixtures of HeLa S3 and HEK293T cell lines. A.) tSNE plot (as in Figure 2d), with cells colored by fraction of reads harboring HeLa S3 specific SNVs relative to hg19 assembly. B.) tSNE using digital gene expression matrices constructed from intronic reads only. Cells are colored by the population of cells from which they derived, with pure HEK293T in red, pure HeLa S3 in yellow, and mixed HEK293T+HeLa S3 in pink

**Figure S4.**
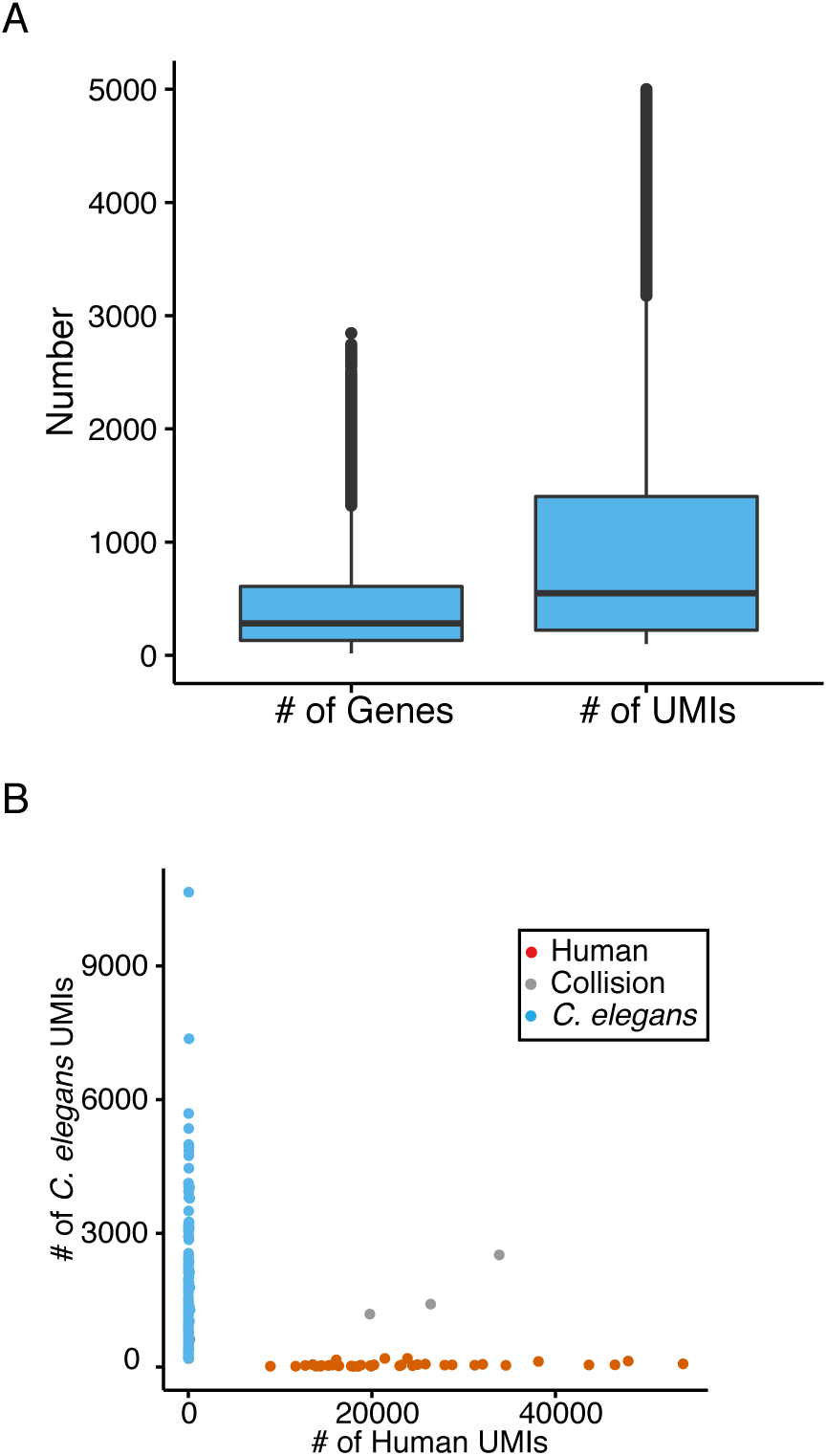
Quality control metrics for *C. elegans* sci-RNA-seq experiments. A.) Distribution of number of protein coding genes and UMI counts (mapping to protein coding genes) detected per *C. elegans* cell. B.) Scatter plot of unique human and *C. elegans* cell UMI counts from a sci-RNA-seq experiment performed on mixture of HEK293T (human) and *C. elegans* cells.

**Table S1.** Summary of experiments

| Experiment ID | Technique version | # of first round barcodes | Cell populations barcoded (# of wells during first round of barcoding) | # of second round barcodes | cells sorted per well |
| --- | --- | --- | --- | --- | --- |
| 1 | Initial protocol | 384 | Pure HEK293T cells (32)<br>Pure HeLa S3 cells (32)<br>Pure NIH/3T3 cells (32)<br>HEK293T & NIH/3T3 cells (96)<br>HeLa S3 & NIH/3T3 cells (96)<br>HEK293T & HeLa S3 cells (96) | 384 | 50 |
| 2 | Optimized protocol | 96 | Pure HEK293T cells (8)<br>Pure HeLa S3 cells (8)<br>HEK293T & NIH/3T3 cells (24)<br>HEK293T & HeLa S3 cells (24)<br>HEK293T & NIH/3T3 nuclei (24) | 96 | 12 |
| 3 | Optimized protocol | 576 | *C. elegans* & HEK293T cells (576) | 960 | 96% of wells: 140 *C. elegans* cells<br>4% of wells: 140 *C. elegans* cells + 10 HEK293T cells |

**Table S2.** Marker genes for cell types identified in sci-RNA-seq *C. elegans* data. A cell is assigned to a cell type (i.e. used when constructing a consensus expression profile for that cell type) if it is in a t-SNE cluster enriched for expression of marker genes listed for that cell type and the individual cell expresses at least one of those marker genes (or *≥* 2 from a set of marker genes where listed).

| Cell type | # cells assigned | # genes expressed (>2% of cells) | Marker genes | References |
| --- | --- | --- | --- | --- |
| Body wall muscle | 11,389 | 4,321 | $\geq 2$ of *lev-11*, *myo-3*, *ttn-1*, *unc-54* | (42) |
| Intestinal/rectal muscle | 165 | 4,561 | *exp-1*, *glb-26* | (43, 44) |
| Pharyngeal muscle | 451 | 3,332 | *myo-2*, *myo-1*, *myo-5*, *tnt-4* | (45) |
| Pharyngeal/buccal epithelia | 678 | 2,781 | *ajm-1*, *dlg-1*, *sma-1*, *agr-1* + absence of *myo-1/2/5* and *tnt-4* | (46–49) |
| Pharyngeal gland | 238 | 3,472 | *phat-4*, *phat-2*, *phat-6* | (50) |
| Hypodermis (hyp1-12) | 2,646 | 4,428 | *dpy-4*, *dpy-5*, *dpy-13* | (51) |
| Seam cells | 1,231 | 4,842 | *grd-13*, *grd-10*, *grd-3* and absence of *dpy-4/5/13* | (52) |
| Cholinergic neurons | 1,038 | 2,265 | *unc-17*, *cho-1*, *cha-1*, *acr-18*, *acr-15* | (53) |
| Ciliated sensory neurons (excluding dopaminergic neurons) | 587 | 2,308 | *R102.2*, *dyf-2*, *che-3*, *nphp-4* | (54) |
| GABAergic neurons | 435 | 1,810 | *unc-25* | (55) |
| Touch receptor neurons | 321 | 1,703 | *mec-17*, *mec-7* | (56) |
| Other interneurons (excluding flp-1(+) interneurons) | 263 | 2,499 | *glr-3* (RIA), *nlp-12* (DVA), Both *des-2* and *deg-3* (PVC/PVD), or *tbh-1* (RIC) | (57–61) |
| flp-1(+) interneurons | 211 | 1,484 | *flp-1* | (62) |
| Canal associated neurons | 187 | 2,241 | $\geq 2$ of *acy-2*, *ace-3*, *mig-6*, *cwn-1* | (63–66) |
| Pharyngeal neurons | 173 | 1,975 | *ser-7*, *eya-1*, *flr-2*, *pha-4* | (67–69) |
| Oxygen sensory neurons | 173 | 2,043 | Both *mec-1*, *lad-2* | (70–74) |
|  |  |  | (ALN/PLN/SDQ), or<br>*gcy-32*, *gcy-36*,<br>*gcy-37*<br>(AQR/PQR/URX), or<br>*flp-17* (BAG) |  |
| Dopaminergic neurons | 70 | 1,821 | *dat-1*, *cat-2* | (75, 76) |
| Unclassified neurons | 2690 | 1,895 | *egl-21*, *sbt-1*, *ida-1*,<br>*casy-1*, *unc-104*, *unc-41* | (77–82) |
| Amphid/phasmid sheath<br>cells | 338 | 4,143 | *vap-1*, *fig-1* | (83) |
| Socket cells | 236 | 3,192 | *grl-1*, *grl-2*, *grd-15* | (52) |
| Unclassified glia | 394 | 2,983 | *kcc-3*, *ram-5*, *daf-6* | (84–86) |
| Somatic gonad precursors<br>(excluding distal tip cells) | 376 | 5,827 | $\geq 2$ of *lin-12*,<br>*inx-9*, *cle-1*, *dgn-1* | (87–90) |
| Distal tip cells | 128 | 5,030 | Both *mig-6*, *nid-1* | (91, 92) |
| Vulval precursor cells | 377 | 5,069 | $\geq 2$ of *lin-12*, *osm-11*,<br>*let-23* | (87, 93, 94) |
| Sex myoblasts | 311 | 4,466 | *egl-15* | (95) |
| Germline | 4301 | 5,086 | $\geq 2$ of *pgl-1*, *cgh-1*,<br>*gld-1*, *iff-1* | (96–99) |
| Rectal epithelial cells | 168 | 4,490 | *grd-1*, *grd-12* | (52, 100) |
| Excretory canal cell | 37 | 5,057 | Both of: *nhr-31*, *ifa-4* | (101, 102) |
| Coelomocytes | 1232 | 1,991 | *cup-4*, *lgc-26*, *unc-122* | (103, 104) |
| Unassigned | 11192 | NA | NA | NA |

